# Mycelial differentiation linked avermectin production in *Streptomyces avermitilis* studied with Raman imaging

**DOI:** 10.1101/2022.09.30.508653

**Authors:** Shumpei Horii, Ashok Zachariah Samuel, Takuji Nakashima, Akira Take, Atsuko Matsumoto, Yoko Takahashi, Masahiro Ando, Haruko Takeyama

## Abstract

*Streptomyces avermitilis* is a gram-positive bacterium that undergoes complex physiological and morphological differentiation during its life cycle, which has implications in secondary metabolites production. Avermectin, produced by *S. avermitilis*, is widely used as an anthelmintic and insecticidal agent. In this study, we have applied Raman microspectroscopic imaging to elucidate the correlation between production of avermectin and the morphological differentiation in *S. avermitilis*. We demonstrate distinctive variations in the localization of avermectin at various morphological stages, such as, substrate mycelium, spore-bearing mycelium, spiral spore chains under solid culture conditions. Under liquid culture condition, however, avermectin is detected only in mycelia after early MII stage of differentiation. Morphological differentiation was observed in liquid and solid cultures, but the chemical profiles of the mycelia were substantially different. Spherical bodies containing avermectin with characteristically different chemical composition to that of spores were also observed under solid culture, which suggests possible release of extracellular vesicles (EVs).

**Key points:**

- Avermectin production is regulated during mycelial differentiation
- Liquid and solid culture conditions affects mycelial differentiation
- Raman microspectroscopic analysis reveals localization profiles of avermectin

## Introduction

Several valuable secondary metabolites with unique properties have been isolated from different microorganisms (Katz et al. 2016). Among them, the filamentous bacterial genus *Streptomyces* contributes to about 70-80% of useful antibiotics, which have several pharmaceutical and agrochemical applications (Bérdy 2005). The mechanism of biosynthesis of secondary metabolites in *Streptomyces* has been investigated in earlier studies (Lee et al. 2021). The stepwise biosynthesis mechanism involves common biomolecular intermediates, such as, amino acids, sugars, fatty acids, and terpenes (Ōmura et al. 2001). Further, the extent of secondary metabolite production is apparently different at various stages of mycelial differentiation (Manteca et al. 2010; Yagüe et al. 2014) Generally, during the life cycle of *Streptomyces* (e.g., *S. coelicolor*) under the solid culture conditions, the seeded spores germinate by growing compartmentalized mycelia (MI) into the agar medium. This initial growth is followed by programmed cell death (PCD) of some parts of the mycelia, but a portion of the surviving mycelial regions grow further as multinucleated mycelia (early MII). Later, this early MII mycelium begins to extend into air forming the aerial mycelia (late MII). Towards the end of this life cycle, a second PCD occurs and most of the surviving late-MII aerial mycelia form spores (Manteca et al. 2010). Similarly, all the above stages of differentiation and growth also occur under liquid culture condition, except for the late MII stage. In most *Streptomyces* species, aerial mycelium formation and sporulation are absent under liquid culture condition, and the growing mycelia form pellets/clumps (Manteca et al. 2008). Importantly, such physiological and morphological differentiation of the parent mycelium has implications in the production of secondary metabolites (Flärdh et al. 2009; McCormick et al. 2012). Gene knockout and proteomic studies have revealed the role of specific microbial hormones in the regulation of secondary metabolism and differentiation in *Streptomyces* (Horinouchi and Beppu 2007). Importantly, the genes involved in secondary metabolism are expressed during MII stage in both solid and liquid culture (Manteca et al. 2010; Yagüe et al. 2014). For instance, a hormone-like signaling molecule called A-factor induces morphological differentiation and regulates streptomycin production in *S. griseus* (Horinouchi 2002). Direct probing of avermectin in various morphologically differentiated regions of *Streptomyces avermitilis*, at single cell level, can help consolidate and validate currently available genomic, proteomic, and morphologic information.

Fluorescent protein labeling is one of the strategies to examine specific protein expression and fluorescence microscopy can reveal their spatial distribution in cells. For instance, localization of carboxylesterase-2, which is involved in biosynthesis of various endogenous and exogenous metabolites, in endoplasmic reticulum has been demonstrated using ratiometric fluorescent probe (Feng et al. 2019). However, this approach cannot reveal the subcellular localization of secondary metabolites directly. On the other hand, time-of-flight secondary ion mass spectrometry (TOF-SIMS) imaging directly revealed the distribution of two antibiotics, undecyl prodigiosin and actinorhodins, on the cell surface of *S. coelicolor* (Vaidyanathan et al. 2008). However, sample destruction is one of the main disadvantages of TOF-SIMS (Spengler 2015).

Raman spectroscopy is a vibrational spectroscopy technique with molecular sensitivity. Raman spectra obtained in single cell imaging analysis can be deconstructed to estimate the relative composition and spatial distribution of multiple chemical constituents simultaneously. (Horii et al. 2020; Samuel et al. 2020) Raman spectroscopy has optical resolution for subcellular investigation, chemical specificity for direct intracellular detection of secondary metabolites, and it is nondestructive. In our earlier studies, we have demonstrated application of Raman microspectroscopy in simultaneously analyzing subcellular localization of several biomolecules including secondary metabolites in *Streptomyces nodosus, Penicillium chrysogenum*, and *Entotheonella*. (Miyaoka et al. 2014; Horii et al. 2020; Samuel et al. 2022; Kogawa et al. 2022). In the present study, using Raman imaging we demonstrate presence of avermectin at specific morphologically differentiated locations of filamentous mycelia of *Streptomyces avermitilis*.

## Materials and methods

### Culturivation of *Streptomyces avermitilis* MA-4680^T^

In this study, *Streptomyces avermectinius* MA-4680^T^ (*S. avermitilis* MA-4680^T^) (Takahashi et al. 2002) was used for Raman microspectroscopic analysis. For liquid culture, a loop of spores of strain MA-4680^T^ was disseminated in 5 mL of seed medium, comprising 2% lactose (FUJIFILM Wako Pure Chemical Corp., Tokyo, Japan), 1.5% malt extract (FUJIFILM Wako Pure Chemical Corp.), 0.5% yeast extract (FUJIFILM Wako Pure Chemical Corp., Tokyo, Japan), and 0.5% dry yeast (ORIENTAL YEAST Corp., Tokyo, Japan). The seed culture was precultured with shaking (200 rpm) at 27 °C. After 2 days of growth, 1 mL of culture was disseminated in 100 mL main medium, comprising 4.5% glucose (FUJIFILM Wako Pure Chemical Corp., Tokyo, Japan), 2.4% peptonized milk (COSMO BIO Corp., Tokyo, Japan), 0.25% yeast extract, 0.25% polypropylene glycol (FUJIFILM Wako Pure Chemical Corp., Tokyo, Japan), and 0.25% dry yeast (adjusted to pH 7.0 before sterilization). The liquid cell culture was incubated with shaking (200 rpm) at 27 °C for 9 days to produce avermectin. For the solid culture, a loop of spores of strain MA-4680^T^ was inoculated in ISP-3 agar medium, comprising 2% oatmeal (Nisshin Seifun Group., Tokyo, Japan), 1.8% agar (FUJIFILM Wako Pure Chemical Corp., Tokyo, Japan), 0.01% FeSO_4_ · 7H_2_O, 0.01% MnCl_2_ · 4H_2_O, and 0.01% ZnSO_4_ · 7H_2_O (adjusted to pH 7.0 before sterilization). The cell culture was incubated at 27 °C for 5 days to form a spore chain.

### LC-ESI-MS and HPLC analyses for avermectin

Avermectin production using strain MA-4680^T^ was measured using an LC/MS spectra (AB Sciex TripleTOF 5600+ System, AB Sciex, Framingham, MA, USA) and HPLC analysis (Hitachi High-Technologies Corp., Tokyo, Japan) (Fig. S1). LC/MS analysis was performed using CAPCELL CORE C18 (3.0 × 100 mm, Osaka Soda Co. Ltd., Osaka, Japan) at 40 °C. For gradient elution, solvent A was ultrapure water with 2 mM ammonium acetate and solvent B was methanol with 2 mM ammonium acetate. Gradient elution was performed with 5% B from 0 to 2 min and 5-100% B from 2 to 12 min. The flow rate was 0.5 mL/min, and injection volume was 2 μL. UV detection was performed using a photodiode array detector. ESI-MS was recorded in the m/z region from 100 to 2000 Da for 12 min. MS analysis was performed using high-resolution ESI-MS (R ≥ 30,000; tolerance for mass accuracy = 5 ppm). HPLC analysis was performed on an Ascentis Express C 18 column (4.6 × 250 mm, Sigma-Aldrich Corp, St Louis, USA) at 40 °C under liner solvent conditions (75:25% CH_3_CN–H_2_O with 1% formic acid for 30 min; flow rate, 1 mL/min; injection volume, 5 μL). UV detection was performed using a photodiode array detector. The standard curve was prepared using serial dilutions of avermectin B1a dissolved in MeOH.

### Raman microspectroscopy and imaging

All Raman spectroscopic measurements were performed using a laboratory-built confocal Raman microspectroscopy setup (Horii et al. 2020). A 532-nm laser was used for excitation. The laser beam was focused onto sample mounted on an inverted microscope using an objective lens (100×, 1.4-NA). The back-scattered Raman light was measured using a spectrometer (spectral resolution of 3.0 cm^−1^). Cell measurements were performed as described below. In the liquid culture condition, the culture broths of strain MA-4680^T^ were sampled at 24h and 168h of cultivation. The culture broths were recovered, then separated into the mycelium and culture-broth using a 40 μm mesh filter. Mycelia trapped on the filter were transferred onto a cover glass. Raman imaging (20 x 20 μm) was conducted with a 0.33-μm step size and 1 s acquisition time. Laser time at the sample was ~4 mW. In the case of solid cultures, 1 cm^3^ of agar culture was collected for Raman mapping measurements. Raman images of mycelia (10 x 10 μm) on the colony surface were recorded with a 0.25-μm step size, 1-s acquisition time (Laser power 4 mW). Melanin pigment produced by the cell greatly affected to the Raman spectrum as background signal on the 5^th^ day of the solid culture. Hence these samples were washed with PBS before the Raman imaging analysis for the removal of melanin pigment. To measure the avermectin B1a reference spectra, commercially available avermectin B1a powder (Fujifilm Wako Pure Chemical Co.) was used. Raman spectral measurements were conducted with 15 mW laser power and 10 s acquisition time.

### Data analysis

Measurement data were preprocessed using IGOR Pro software (WaveMetrics, Inc., Lake Oswego, OR, USA). All spectra acquired from the Raman mapping measurements were combined into a single matrix. Intensity correction was performed using the halogen lamp spectrum and wavenumber calibration was performed using the Raman spectrum of indene. Before MCR-ALS calculation, noise reduction was performed using singular value decomposition (SVD). MCR-ALS was performed as previously described (Lee and Seung 1999; Azzouz and Tauler 2008; Ando and Hamaguchi 2013; Horii et al. 2020; Samuel et al. 2021).

## Results

### Avermectin production in *S. avermitilis* MA-4680^T^ under liquid culture

The mycelial pellets of *S. avermitilis* MA-4680^T^ collected after 24h and 168h cultivation in liquid medium were analyzed using Raman microscpectroscopic imaging. Raman spectral components corresponding to four different biomolecules (Fig. 1a), including that of secondary metabolite avermectin, were extracted from the whole Raman image data using MCR-ALS multivariate analysis. The Raman spectrum of component 1 represents proteins: its features include peaks at 1004 cm^−1^, the ring breathing mode of phenylalanine and tryptophan residues; 1270 cm^−1^, Amide III; 1449 cm^−1^, C–H bending; and 1654 cm^−1^, Amide I (Huang et al. 2004). The component 2, with Raman bands at 746, 1126, 1304, 1335, and 1580 cm^−1^, represents cytochrome *b* (Okada et al. 2012; Kakita et al. 2013). Component 3 shows Raman bands associated with lipids: 1080 cm^−1^, ν(C–C) stretching vibrations; 1304 cm^−1^, δ(CH_2_) twisting vibrations; and 1440 cm^−1^, δ(CH_2_) wagging vibrations. Additionally, Raman band derived from C=C stretching vibration, which represents the degree of unsaturation of fatty acids, was observed at 1657 cm^−1^. The Raman band at 1747 cm^−1^ indicates ester vibration (Samuel et al. 2022). Component 4 shows two prominent Raman bands of avermectin at 1629 cm^−1^ and 1676 cm^−1^ assignable to conjugated (-C=C-C=C-) and non-conjugated trans C=C (Fig. 1c) (Schrader and Meier 1974; Berdyugin et al. 1988; Meurens et al. 2005).

**Fig. 1.**
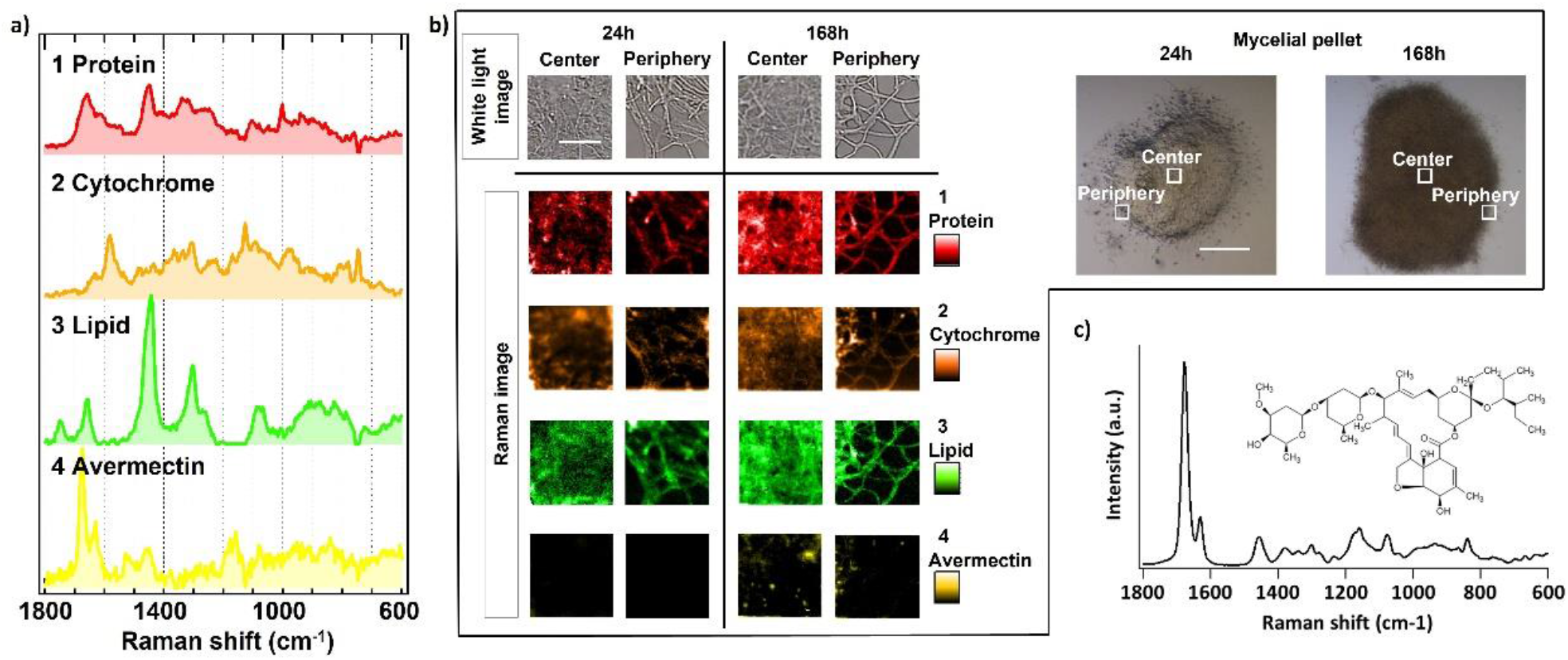
Raman microspectroscopic analysis of molecular composition of mycelia produced under liquid culture conditions. MCR-ALS resolved Raman spectra (a) corresponding to (1) protein, (2) cytochrome *b*, (3) lipid, and (4) avermectin and (b) spatial distribution maps (scale bar 5 μm) for components 1–4 together with corresponding optical microscopy images (Two photos of whole mycelial pellets are shown on the right; scale bar 0.1 mm). A full list is provided in the Figure S4. (c) Raman spectrum of avermectin B1 powder sample.

The intracellular molecular distribution images corresponding to spectral components given in Fig. 1a are shown in Fig. 1b. The protein, cytochrome *b*, lipid, and avermectin can be seen distributed in the cytoplasm of the individual mycelia (Fig. 1b – periphery). Avermectin was detected only in one of the samples collected at 168h of incubation and was highly concentrated at certain locations of the center of the mycelial pellet. Interestingly, avermectin distribution is not uniform even in individual mycelia, but avermectin is concentrated at certain regions of the individual mycelia (Fig. S2).

### Avermectin production in *S. avermitilis* MA-4680^T^ under solid culture

*S. avermitilis* MA-4680^T^ was also cultured on agar medium, and Raman imaging was conducted using the samples collected after 1 to 5 days of solid culture (Fig. 2). As explained in introduction section, under solid culture condition late MII mycelial differentiation can be observed. After 72h of incubation, spiral spore chains (SSCs) were observed in the samples. Five Raman spectral components were obtained in the MCR-ALS analysis of the Raman image data (Fig. 2a). Based on the Raman bands MCR spectral components 1, 2, 3, and 4 have been assigned to protein, cytochrome, lipid, and avermectin, respectively (see the previous section for spectrum assignment). Compound 5 has been assigned to carotenoid, which clearly exhibits vibrations of methyl group (C–CH_3_) at 1007 cm^−1^, C-C at 1156 cm^−1^, and C=C at 1524 cm^−1^ (De Oliveira et al. 2010). The photos of substrate mycelium (SM) and SSCs produced after late MII stage mycelial differentiation are shown in Fig. 2b. The corresponding Raman images are shown in Fig. 2c. Proteins, lipids (droplet like features; Kaddor et al. 2009), and cytochrome show similar mycelial distribution to that seen in mycelia grown under liquid culture conditions. However, avermectin is not detected in SM mycelia even after early MII stage differentiation (i. e., after 24 hours; Fig. 2c). Raman images show that the chemical composition of SM and SSCs are characteristically different (Fig. 2c). Carotenoid is rich in SSCs compared to SM. Avermectin is found only in SSCs and not in SM.

**Fig. 2.**
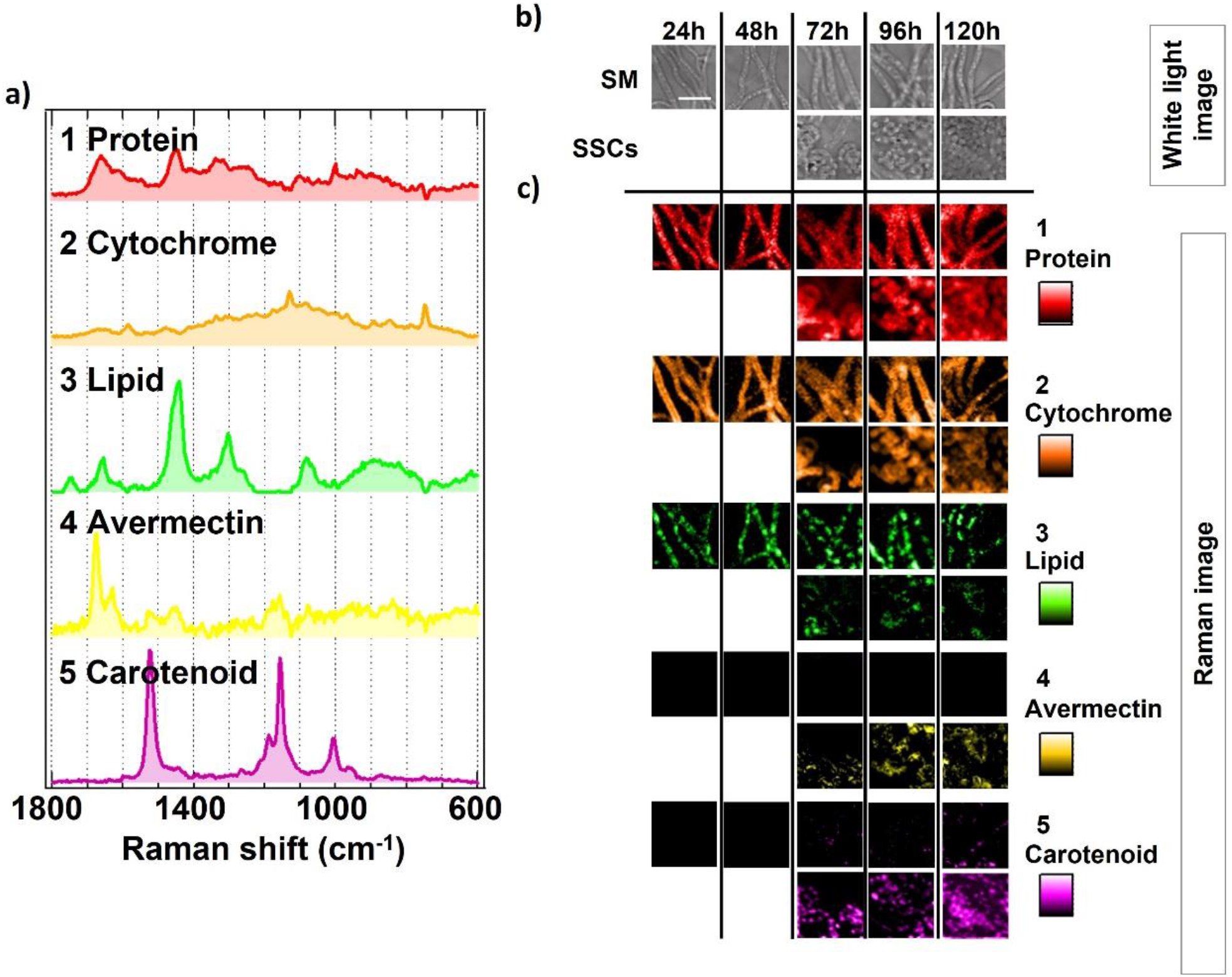
Raman microspectroscopic analysis of molecular composition of various differentiated mycelia produced under solid culture conditions. (a) MCR-ALS resolved Raman spectra corresponding to (1) protein, (2) cytochrome b (3) lipid, (4) avermectin, and (5) carotenoid. A full list is provided in the Figure S4. (b) Optical images of SM and SSCs collected from solid culture at different incubation times and (c) corresponding Raman images of components 1–5. Scale bar = 5 μm.

Differentiated arial mycelia produced after late MII stage differentiation have SM base, spore-bearing mycelium (SBM), and SSCs. A pictorial depiction of differentiated mycelia is shown in Fig. 3a. To perform detailed chemical analysis, SM, SBM, and SSCs were separated on the 5^th^ day of the solid culture, and each separate domain was analyzed with Raman imaging. In the Raman images, avermectin can be found localized in the SBM and SSCs but not in the SM (Fig. 3b). This reaffirms the results presented in Fig. 2.

**Fig. 3.**
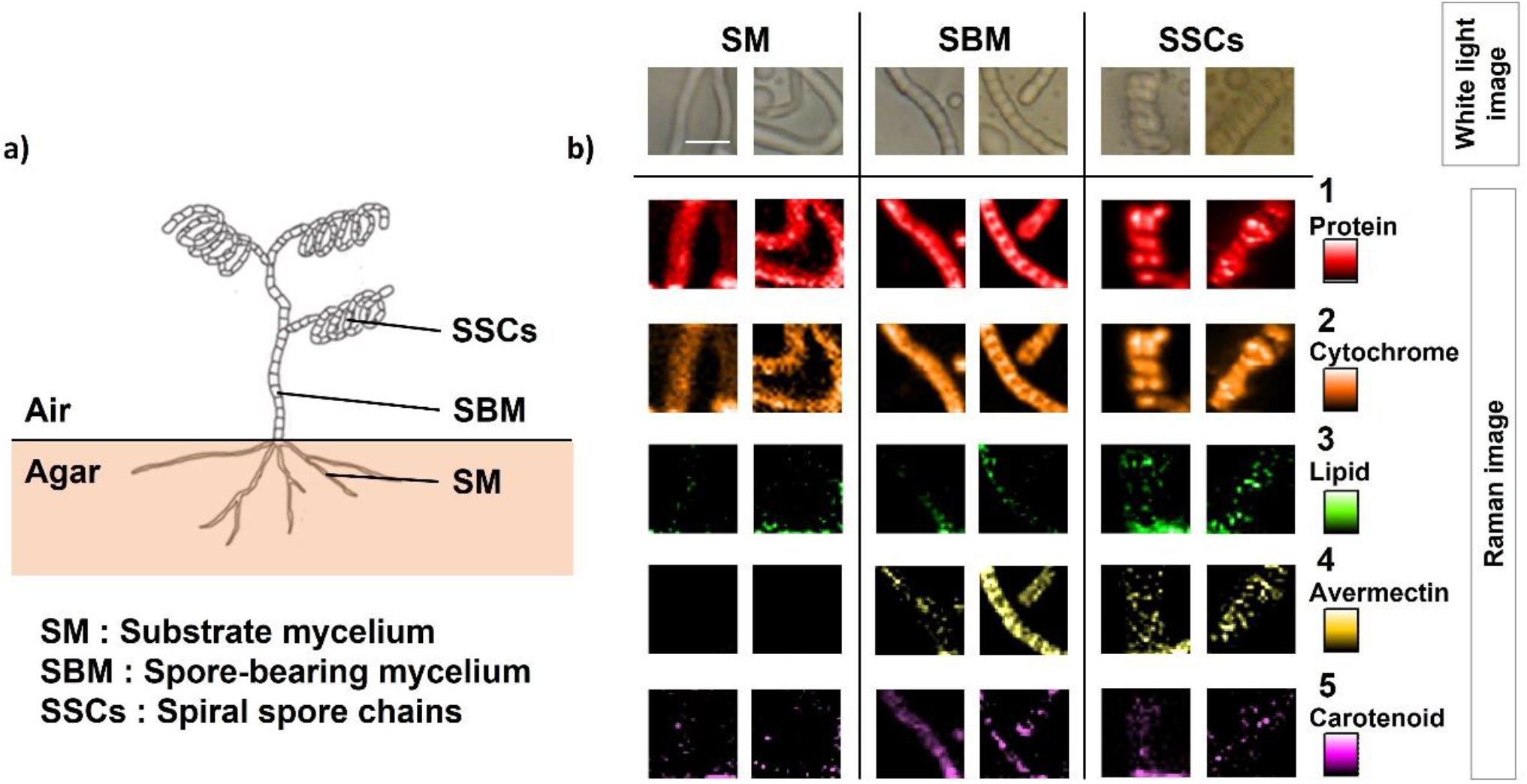
Late MII mycelial differentiation and Raman images of SM, SBM, and SSCs. (a) pictorial depiction of late MII mycelial differentiation and (b) Raman images corresponding to (1) protein, (2) cytochrome b (3) lipid, (4) avermectin, and (5) carotenoid with corresponding optical microscopy images. Scale bar = 2 μm.

After 72h of culture, spherical structures with chemical composition characteristically different from that of spores were also detected in the agar culture of *S. avermitilis* MA-4680^T^. Raman spectral components detected in these structures were assigned to protein, lipid, and avermectin, respectively (Fig. 4, S3). Avermectin was observed in only a fraction of the spherical structures analyzed (Fig. 4). Specifically, 29 spherical structures were measured and avermectin was detected in 19 of them. Importantly, carotenoid was absent in them, which is a unique component observed in SSCs. Distinct chemical composition suggests a different origin of the structures, such as, extracellular vesicles (EVs) released from SM. However, this conjecture is not conclusive.

**Fig. 4.**
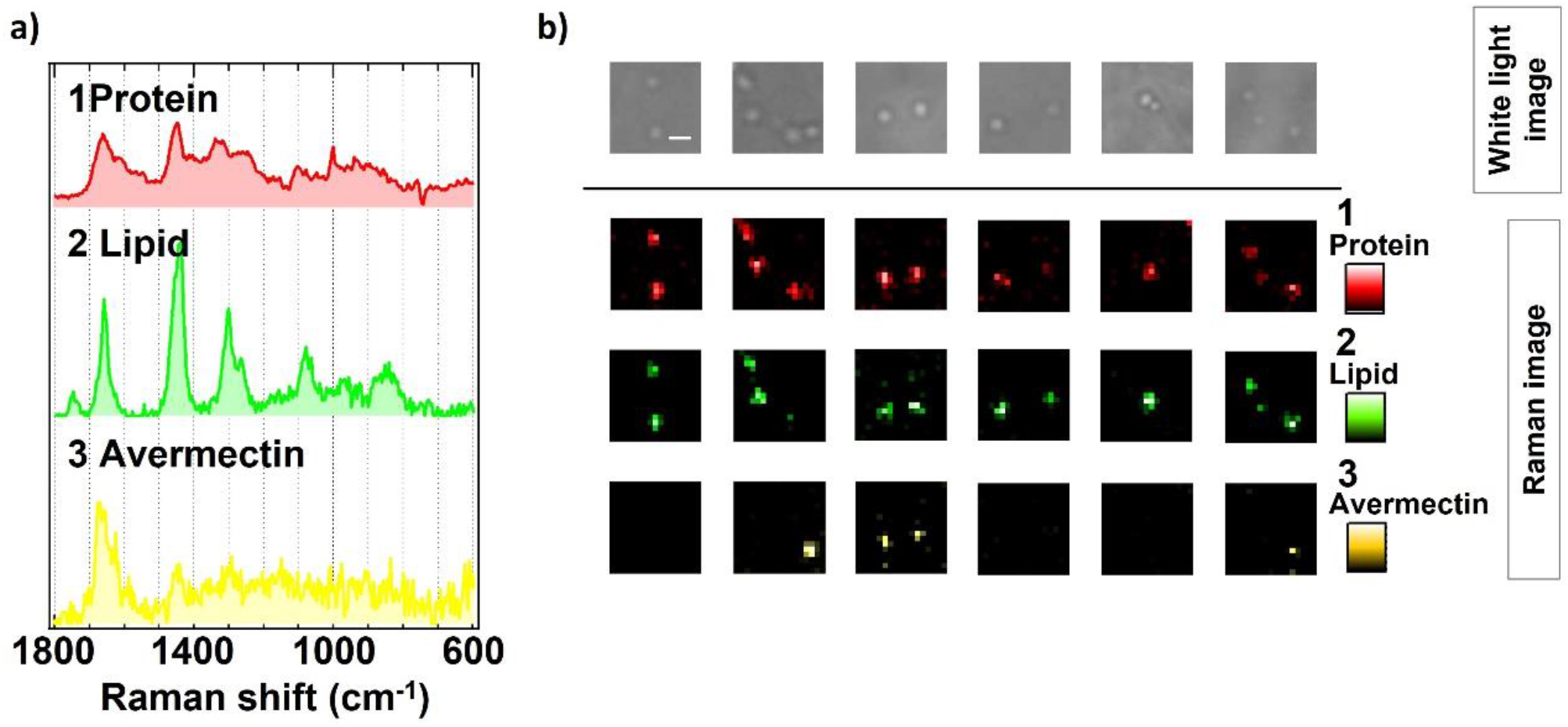
Raman spectra of chemical components detected in the spherical structures observed in solid culture condition and the corresponding Raman images. Scale bar = 1 μm.

## Discussion

*S. avermitilis* MA-4680^T^ produces avermectins, which are a series of 16-membered pentacyclic compounds with a disaccharide of methylated deoxysugar L-oleandrose polyketides (Ikeda and Ōmura 1997). Studies have shown that specific morphological differentiation can activate secondary metabolite production in microbes (Manteca et al. 2010; Yagüe et al. 2014). Using Raman imaging, we have shown that mycelial distribution of proteins, lipids, cytochrome, carotenoid and avermectin differed characteristically in *S. avermitilis* at different stages of mycelial differentiation. Mycelial pellet analyzed at 24h of liquid culture did not show appreciable avermectin, which suggests that the growth is in the MI stage. This is consistent with the observation in *S. coelicolor* (Manteca et al. 2008). However, after 168h of incubation, avermectin was concentrated at the center of the mycelial pellet. This observation is consistent with the differentiation of MI mycelia into early MII mycelia, and subsequent production of avermectin (Manteca et al. 2008). Previously reported proteome analysis revealed that genes/proteins involved in secondary metabolism are expressed/translated during MII stage in *S. coelicolor* (Manteca et al. 2010; Yagüe et al. 2014), which supports our observation. It is also important to notice that avermectin is not uniformly distributed in individual mycelia but concentrated at certain mycelial locations. Such local enrichment along the mycelia has been observed in other *Streptomyces* sp. (Miyaoka et al. 2014).

Under solid culture, late MII mycelia of *S. avermitilis* differentiates into arial mycelia, which allowed us to examine avermectin production in different mycelial domains after this differentiation. We collected the samples from solid culture at different time intervals (24h to 120h), which were progressively at different stages of differentiation. Raman imaging revealed small amount of carotenoid production in substrate mycelia after 72h, which were apparently in the MII stage of differentiation. Carotenoid content continued to increase from 72h to 120h (Fig. 2c). It is important to note that MII differentiated mycelia under liquid culture did not produce carotenoid. Therefore, this change in chemical composition of mycelia suggest beginning of the next stage of mycelial differentiation. Supporting this argument, carotenoid composition was substantially high in the SBM and SSCs, which were produced at the late MII stage of differentiation. More interestingly, this compositional difference also correlates with higher avermectin production in the SBM and SSCs (Fig. 3b). However, unlike in the case of liquid culture, avermectin was not observed in these late MII substrate mycelia. The biosynthetic pathway of avermectin is well established and it mainly involves three stages: 1) formation of early aglycone derived from polyketide, 2) production of avermectin aglycone by modification of early aglycone, and 3) glycosylation of avermectin aglycone. Labeling experiments have shown that the aglycone portion of avermectin is derived from various fatty acids (Ikeda and Ōmura 1997). Thus, the currently accepted mechanism of avermectin production doesn’t involve carotenoid. Therefore, its production is not directly related to avermectin biosynthesis, but it may play role in protecting arial mycelia from environmental stress (Avalos et al. 2015). However, the exact role of carotenoid remains unclear. Production of antibiotics during sporulation, on the other hand, is not unusual. For instance, several antibiotics are produced by *Streptomyces* sp. during sporulation, such as, neomycin by *S. fradiae*, streptomycin by *S. griseus* (Barabas and Szabó 1968), and actinorhodin in *S. coelicolor* (Čihák et al. 2017). Antibiotics can protect spores against environmental factors prior to reactivation from the dormant stage (Barabas and Szabó 1968).

Absence of avermectin in MII mycelia (substrate mycelia, SM) is another surprisingly distinct observation in solid culture compared to the liquid culture. The gene expression involved in secondary metabolite biosynthesis (86% of all transcripts) is found to be similar in *Streptomyces* sp. under both liquid and solid culture conditions (Yagüe et al. 2014). However, according to Yagüe et al., the expression of genes related to secondary metabolite biosynthesis (e.g., coelichelin and desferrioxamine) were upregulated in liquid culture conditions. Based on our results, we believe that avermectin production is also similarly upregulated under liquid culture (down regulated under solid culture).

The release of extracellular vesicles is a widespread and important phenomenon in bacteria (Faddetta et al. 2022). Extracellular vesicles have characteristics that can mediate the transfer of DNA fragments, autolysins, cytotoxins, virulence factors, and various other biomolecules (Alaniz et al. 2007; Furuta et al. 2009; Biller et al. 2014; Fulsundar et al. 2014) These secreted products help bacteria-bacteria and bacteria-host interactions (Kim et al. 2015). *S. coelicolor* is known to produce membrane vesicles, which include amino acids and their precursors, vitamins, components of purine and pyrimidine metabolism, components of carbon metabolism, and antibiotics (Schrempf et al. 2011). In our study, compositionally different but avermectin containing spherical structures were observed under solid culture condition (Fig. 4). The possibility of this being EVs extruded from the MII mycelia of *S. avermitilis* cannot be ruled out. However, such large EVs-like (~1 μm) structures were not observed in our liquid culture experiments.

We have demonstrated an application of Raman imaging in chemical profiling of microbes during mycelial differentiation. We demonstrated that the chemical constitution of mycelia changes considerably during morphological differentiation. MII stage morphological differentiation is observed in liquid and solid cultures, but the chemical profiles of the mycelia were substantially different. Hence, morphological differentiation alone is not a good indicator of metabolic regulation in microbes. Analysis of chemical composition as well as subcellular localization information is necessary for delineating intricate biosynthetic mechanism involved. We believe that Raman imaging has several such potential applications in investigating secondary metabolite production/regulation in microbes. Understanding detailed chemical evolution during the lifecycle of microbes can help optimizing culture conditions for effective antibiotic production and might also lead to the discovery of novel antibiotics.

## Data Availability

The datasets generated during and/or analyzed during the current study are available from the corresponding author on reasonable request.

## Supporting information

The online version contains supplementary materials.

## Acknowledgements

This work was supported by a Grant-in-Aid for Scientific Research S (no. 17H06158).

## Author contributions

SH, MA, TN, AZS, AT, AM, YT and HT conceived the idea and designed the experiments. SH, TN and AT conducted LC/MS and HPLC. SH conducted Raman spectroscopic measurements and MCR data analysis. SH and AZS performed data analysis, interpretation, and manuscript writing. Software code written by MA was used in the data analysis. HT, MA and TN also contributed to manuscript writing and discussions.

## Statements and Declarations

### Ethics approval

This article does not contain any study with human participants or animals performed by any of the authors.

## REFERENCES

Alaniz RC, Deatherage, BL, Lara JC, Cookson BT (2007) Membrane vesicles are immunogenic facsimiles of Salmonella typhimurium that potently activate dendritic cells, prime B and T cell responses, and stimulate protective immunity in vivo. J Immunol 179:7692–7701. doi: 10.4049/jimmunol.179.11.7692

Ando M, Hamaguchi H (2013) Molecular component distribution imaging of living cells by multivariate curve resolution analysis of space-resolved Raman spectra. J Biomed Opt 19:011016. doi: 0.1117/1.JBO.19.1.011016

Avalos J, Carmen Limón M (2015) Biological roles of fungal carotenoids. Curr Genet 61:309–324. doi: 10.1007/s00294-014-0454-x

Azzouz T, Tauler R (2008) Application of multivariate curve resolution alternating least squares (MCR-ALS) to the quantitative analysis of pharmaceutical and agricultural samples. Talanta 74:1201–1210. doi: 10.1016/j.talanta.2007.08.024

Barabás G, Szabó G (1968) Role of streptomycin in the life of Streptomyces griseus: streptidine-containing fractions in the cell walls of *Streptomyces griseus* strains. Can J Microbiol 14:1325–1331. doi: 10.1139/m68-222

Bérdy J (2005) Bioactive microbial metabolites. J Antibiot 58:1–26. doi: 10.1038/ja.2005.1

Berdyugin VV, Burshtein KY, Anikin NA, Krasnaya ZA, Stytsenko TS (1988) Resonance raman light scattering spectra and cross section and the electronic structure of polyene compounds. Russ Chem Bull 37:1378–1382. doi: 10.1007/BF00962744

Biller SJ, Schubotz F, Roggensack SE, Thompson AW, Summons RE, Chisholm SW (2014) Bacterial vesicles in marine ecosystems. Science 343:183–186. doi: 10.1126/science.1243457

Čihák M, Kameník Z, Šmídová K, Bergman N, Benada O, Kofroňová O, Petříčková K, Bobek J (2017) Secondary metabolites produced during the germination of *Streptomyces coelicolor*. Front Microbiol 8:2495. doi: 10.3389/fmicb.2017.02495

De Oliveira VE, Castro HV, Edwards HGM, de Oliveira LF (2010) Carotenes and carotenoids in natural biological samples: a Raman spectroscopic analysis. J Raman Spectrosc 41:642–650. doi: 10.1002/jrs.2493

Faddetta T, Renzone G, Vassallo A, Rimini E, Nasillo G, Buscarino G, Agnello S, Licciardi M, Botta L, Scaloni A, Palumbo Piccionello A, Puglia AM, Gallo G (2022) *Streptomyces coelicolor* vesicles: many molecules to be delivered. Appl Environ Microbiol 88:e0188121. doi: 10.1128/AEM.01881-21

Feng L, Ning J, Tian X, Wang C, Zhang L, Ma X, James TD (2019) Fluorescent probes for bioactive detection and imaging of phase II metabolic enzymes. Coord Chem Rev 399:213026. doi: 10.1016/j.ccr.2019.213026

Flärdh K, Buttner MJ (2009) *Streptomyce*s morphogenetics: dissecting differentiation in a filamentous bacterium. Nat Rev Microbiol 7:36–49. doi: 10.1038/nrmicro1968

Fulsundar S, Harms K, Flaten GE, Johnsen PJ, Chopade BA, Nielsen KM (2014) Gene transfer potential of outer membrane vesicles of Acinetobacter baylyi and effects of stress on vesiculation. Appl Environ Microbiol 80:3469–3483. doi: 10.1128/AEM.04248-13

Furuta N, Tsuda K, Omori H, Yoshimori T, Yoshimura F, Amano A (2009) Porphyromonas gingivalis outer membrane vesicles enter human epithelial cells via an endocytic pathway and are sorted to lysosomal compartments. Infect Immun 77:4187–4196. doi: 10.1128/IAI.00009-09

Horii S, Ando M, Samuel AZ, Take A, Nakashima T, Matsumoto A, Takahashi Y, Takeyama H (2020) Detection of penicillin G produced by *Penicillium chrysogenum* with Raman microspectroscopy and multivariate curve resolution-alternating least-squares methods. J Nat Prod 83:3223–3229. doi: 10.1021/acs.jnatprod.0c00214

Horinouchi S (2002) A microbial hormone, A-factor, as a master switch for morphological differentiation and secondary metabolism in *Streptomyces griseus*. Front Biosci 7:2045–2057. doi: 10.2741/A897

Horinouchi S, Beppu T (2007) Hormonal control by A-factor of morphological development and secondary metabolism in *Streptomyces*. Proc Jpn Acad Ser B Phys Biol Sci 83:277–295. doi: 10.2183/pjab/83.277

Huang WE, Griffiths RI, Thompson IP, Bailey MJ, Whiteley AS (2004) Raman microscopic analysis of single microbial cells. Anal Chem 76:4452–4458. doi: 10.1021/ac049753k

Ikeda H, Ōmura S (1997) Avermectin Biosynthesis. Chem Rev 97:2591–2610. doi: 10.1021/cr960023p

Kaddor C, Biermann K, Kalscheuer R, Steinbüchel A (2009) Analysis of neutral lipid biosynthesis in *Streptomyces avermitilis* MA-4680 and characterization of an acyltransferase involved herein. Appl Microbiol Biotechnol 84:143–155. doi: 10.1007/s00253-009-2018-4

Kakita M, Okuno M, Hamaguchi HO (2013) Quantitative analysis of the redox states of cytochromes in a living L929 (NCTC) cell by resonance Raman microspectroscopy. J Biophotonics 6:256–259. doi: 10.1002/jbio.201200042

Katz L, Baltz RH (2016) Natural product discovery: past, present, and future. J Ind Microbiol Biotechnol 43:155–176. doi: 10.1007/s10295-015-1723-5

Kim JH, Lee J, Park J, Gho YS (2015) Gram-negative and Gram-positive bacterial extracellular vesicles. Semin Cell Dev Biol 40:97–104. doi: 10.1016/j.semcdb.2015.02.006

Kogawa M, Miyaoka R, Hemmerling F, Ando M, Yura K, Ide K, Nishikawa Y, Hosokawa M, Ise Y, Cahn JKB, Takada K, Matsunaga S, Mori T, Piel J, Takeyama H (2022) Single-cell metabolite detection and genomics reveals uncultivated talented producer. PNAS Nexus 1:pgab007. doi: 10.1093/pnasnexus/pgab007

Lee DD, Seung HS (1999) Learning the parts of objects by non-negative matrix factorization. Nature 401:788–791. doi: 10.1038/44565

Lee N, Hwang S, Kim W, Lee Y, Kim JH, Cho S, Kim HU, Yoon YJ, Oh MK, Palsson BO, Cho BK (2021) Systems and synthetic biology to elucidate secondary metabolite biosynthetic gene clusters encoded in *Streptomyces* genomes. Nat Prod Rep 38:1330–1361. doi: 10.1039/d0np00071j

Manteca A, Alvarez R, Salazar N, Yagüe P, Sanchez J (2008) Mycelium differentiation and antibiotic production in submerged cultures of Streptomyces coelicolor. Appl Environ Microbiol 74:3877–3886. doi: 10.1128/AEM.02715-07

Manteca A, Jung HR, Schwammle V, Jensen ON, Sanchez J (2010) Quantitative proteome analysis of Streptomyces coelicolor nonsporulating liquid cultures demonstrates a complex differentiation process comparable to that occurring in sporulating solid cultures. J Proteome Res 9:4801–4811. doi: 10.1021/pr100513p

McCormick JR, Flärdh K (2012) Signals and regulators that govern *Streptomyces* development. FEMS Microbiol Rev 36:206–231. doi: 10.1111/j.1574-6976.2011.00317.x

Meurens M, Baeten V, Yan SH, Mignolet E, Larondelle Y (2005) Determination of the conjugated linoleic acids in cow’s milk fat by Fourier transform Raman spectroscopy. J Agric Food Chem 53:5831–5835. doi: 10.1021/jf0480795

Miyaoka R, Hosokawa M, Ando M, Mori T, Hamaguchi HO, Takeyama H. (2014) In situ detection of antibiotic amphotericin B produced in *Streptomyces nodosus* using Raman microspectroscopy. Mar Drugs 12:2827–2839. doi: 10.3390/md12052827

Okada M, Smith NI, Palonpon AF, Endo H, Kawata S, Sodeoka M, Fujita K (2012) Label-free Raman observation of cytochrome *c* dynamics during apoptosis. PNAS 109:28–32. doi: 10.1073/pnas.1107524108

Ōmura S, Ikeda H, Ishikawa J, Hanamoto A, Takahashi C, Shinose M, Takahashi Y, Horikawa H, Nakazawa H, Osonoe T, Kikuchi H, Shiba T, Sakaki Y, Hattori M (2001) Genome sequence of an industrial microorganism *Streptomyces avermitilis*: deducing the ability of producing secondary metabolites. PNAS 98:12215–12220. doi: 10.1073/pnas.211433198

Samuel AZ, Miyaoka R, Ando M, Gaebler A, Thiele C, Takeyama H (2020) Molecular profiling of lipid droplets inside HuH7 cells with Raman micro-spectroscopy. Commun Biol 3:1–10. doi: 10.1038/s42003-020-1100-4

Samuel AZ, Horii S, Ando M, Takeyama H (2021) Deconstruction of Obscure Features in SVD-Decomposed Raman Images from P. chrysogenum Reveals Complex Mixing of Spectra from Five Cellular Constituents. Anal Chem 93:12139–12146. doi: 10.1021/acs.analchem.1c02942

Samuel AZ, Horii S, Nakashima T, Shibata N, Ando M, Takeyama H (2022) Raman Microspectroscopy Imaging Analysis of Extracellular Vesicles Biogenesis by Filamentous Fungus Penicilium chrysogenum. Adv Biol 2101322. doi: 10.1002/adbi.202101322

Schrader B, Meier W (1974) Raman/infrared atlas of organic compounds. VCH.

Schrempf H, Koebsch I, Walter S, Engelhardt H, Meschke H (2011) Extracellular *Streptomyces* vesicles: amphorae for survival and defence. Microb Biotechnol 4:286–299. doi: 10.1111/j.1751-7915.2011.00251.x

Spengler B (2015) Mass spectrometry imaging of biomolecular information. Anal Chem 87:64–82. doi: 10.1021/ac504543v

Takahashi Y, Matsumoto A, Seino A, Ueno J, Iwai Y, Ōmura S (2002) *Streptomyces avermectinius* sp. nov., an avermectin-producing strain. Int J Syst Evol Microbiol 52:2163–2168. doi: 10.1099/00207713-52-6-2163

Vaidyanathan S, Fletcher JS, Goodacre R, Lockyer NP, Micklefield J, Vickerman JC (2008) Subsurface biomolecular imaging of *Streptomyces coelicolor* using secondary ion mass spectrometry. Anal Chem 80:1942–1951. doi: 10.1021/ac701921e

Yagüe P, Rodríguez-García A, López-García MT, Rioseras B, Martín JF, Sánchez J, Manteca A (2014) Transcriptomic analysis of liquid non-sporulating *Streptomyces coelicolor* cultures demonstrates the existence of a complex differentiation comparable to that occurring in solid sporulating cultures. PLoS One 9:e86296. doi: 10.1371/journal.pone.0086296

