## Supplementary Figures for "Mycelial differentiation linked avermectin production in *Streptomyces avermitilis* studied with Raman imaging"

a)

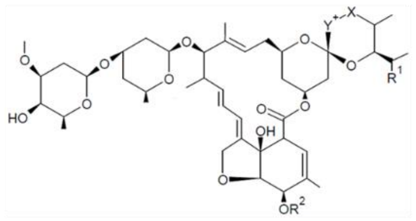

| Avermectin | $R_1$ | $R_2$ | X-Y |
|---|---|---|---|
| A1a | $CH_3CH_2$ | $CH_3$ | CH=CH |
| A1b | $CH_3$ | $CH_3$ | CH=CH |
| A2a | $CH_3CH_2$ | $CH_3$ | CH-CH(OH) |
| A2b | $CH_3$ | $CH_3$ | CH-CH(OH) |
| B1a | $CH_3CH_2$ | H | CH=CH |
| B1b | $CH_3$ | H | CH=CH |
| B2a | $CH_3CH_2$ | H | CH-CH(OH) |
| B2b | $CH_3$ | H | CH-CH(OH) |

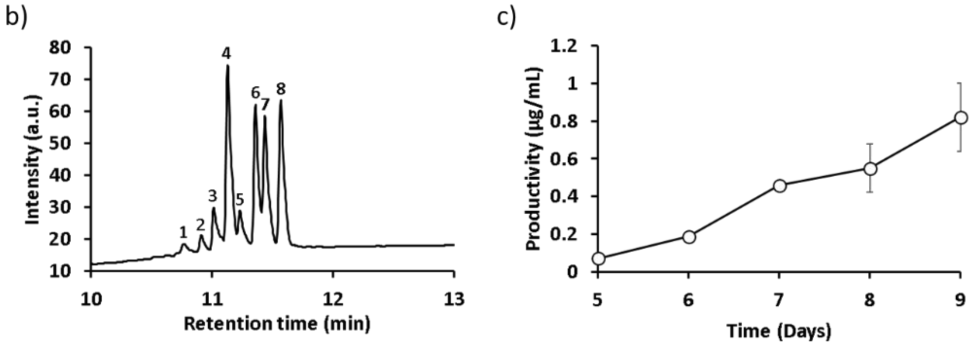


**Fig. S1** Measurement of avermectin produced by *S. avermitilis* MA-4680$^T$ in liquid culture

*S. avermitilis* MA-4680$^T$ is known to produce 11 avermectin analogs. The antibiotic produced by MA-4680$^T$ strain was measured via HPLC and LC/MS. Thus, the production of eight avermectin analogs was confirmed. On each day, quantification of avermectin B1a was performed using the peak area. Productivity of avermectin B1a increased in the 5-day-old liquid culture. a) Structure of avermectin analogs. b) HPLC chromatograms of the culture extracts with MtOH. The peaks were assigned via LC/MS as follows; peak 1: avermectin B2b; peak 2: B2a; peak 3: A2b; peak 4: A2a; peak5: B1b; peak6: B1a; peak7: A1b; peak8: A1a. c) Quantification of avermectin B1a production in liquid culture.

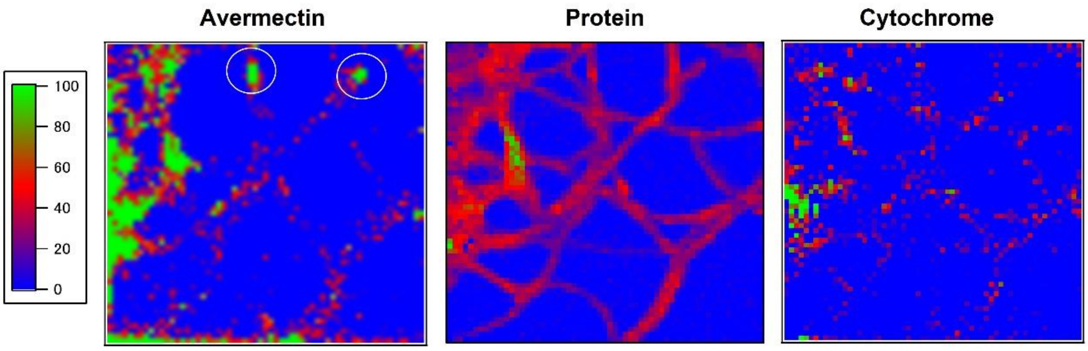


**Fig. S2** Raman images of avermectin, protein and cytochrome b in individual mycelia detected at the periphery of the mycelial pellet obtained in liquid culture at 168h. Avermectin enriched at two mycelial locations are circled. At the same time, protein is distributed equally throughout the cytoplasm as expected.

Cytochrome shows characteristic distribution as expected (Horii et al. 2020).

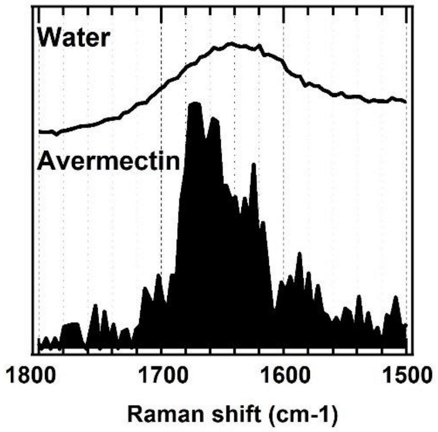


**Fig. S3** A comparison of Avermectin spectrum resolved in the MCR-ALS analysis and water bending Raman band.

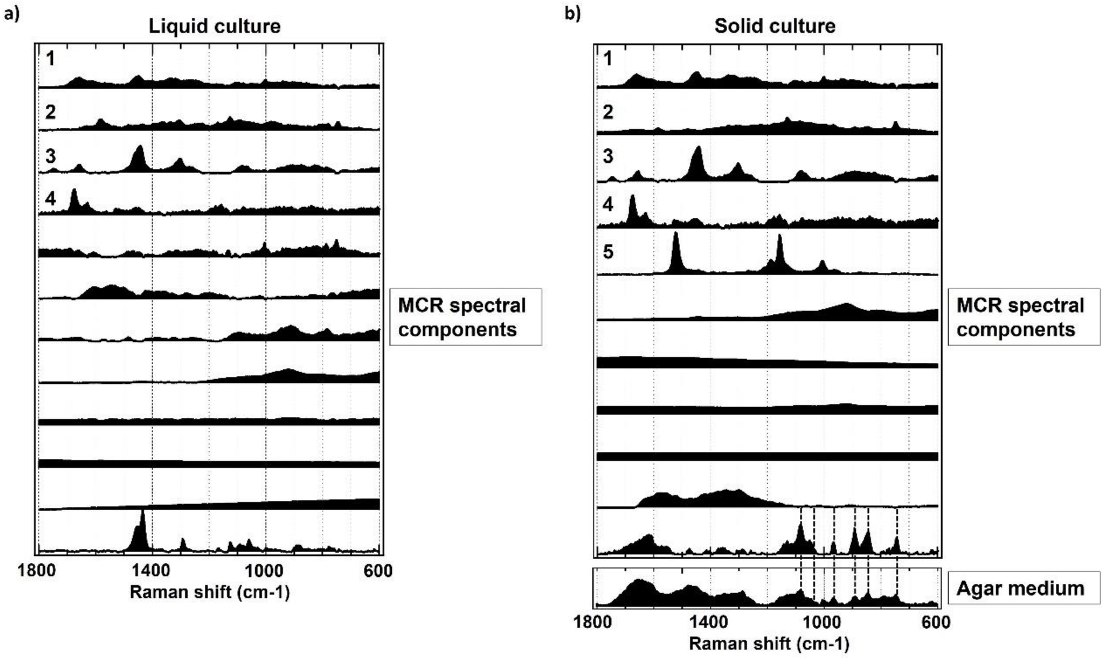


**Fig. S4** Full list of MCR spectral components used in modelling the Raman data from *S. avermitilis*. a) In liquid culture, the top four MCR components have interpretable and relevant spectral components representing biomolecules. Component 8 is the glass background. Component 12 is residual oil/wax from the culture medium found outside mycelial regions (very rare). b) In solid culture, the top five MCR components have interpretable and relevant spectral components representing biomolecules. Component 6 is the glass background. Component 11 has contribution from agar medium. The broad components are treated as background in both cases.
